# Allele-specific expression reveals genetic drivers of tissue regeneration in mice

**DOI:** 10.1101/2022.09.23.509223

**Authors:** Heather E. Talbott, Katya L. Mack, Michelle Griffin, Nicholas J. Guardino, Jennifer B.L. Parker, Amanda F. Spielman, Michael F. Davitt, Shamik Mascharak, Mark J. Berger, Derrick C. Wan, Hunter B. Fraser, Michael T. Longaker

## Abstract

In adult mammals, skin wounds typically heal by scarring rather than through regeneration. In contrast, “super-healer” MRL mice have the unusual ability to regenerate ear punch wounds, yet the molecular basis for this regeneration remains elusive. Here, in hybrid crosses between MRL and non-regenerating mice, we use allele-specific gene expression to identify *cis*-regulatory variation associated with ear regeneration. Analyzing three major wound cell populations, we identified extensive strain- and tissue- specific *cis-*regulatory divergence associated with differences in healing outcomes. Genes with *cis-*regulatory differences specifically in fibroblasts were associated with wound healing phenotypes and pathways, and were enriched near genetic markers associated with ear-healing in a genetic cross. Finally, we demonstrated that one of these genes, *Cfh*, could be applied ectopically to accelerate wound repair and induce regeneration in typically fibrotic wounds. Overall, our results provide insight into the molecular drivers of regeneration in MRL mice with potential clinical implications.

## Introduction

Fibrosis, or the replacement of functional tissue with non-functional connective tissue, can result from tissue damage to any organ in the human body. In the skin, fibrosis occurs as scarring and has major consequences for skin form and function. Scars lack the structures (e.g., hair, glands) of normal skin, compromising skin’s normal barrier system and its ability to thermoregulate, and are weaker and less flexible than uninjured skin. Healing via scarring has major consequences for human health: scarring can cause disfigurement, functional loss, and reduced quality of life (Bayat et al., 2003; desJardins-Park et al., 2021). Despite the substantial clinical burden scars impose, there are no current therapies that induce scar-free healing in humans.

In contrast to humans, some other species possess the ability to regenerate skin after injury without scar formation. The Murphy Roths Large (MRL) mouse represents a valuable biomedical model for studying mammalian wound regeneration. While injuries to mammalian skin and other organs typically heal via formation of fibrotic scar tissue, MRL and its progenitor strain, the Large (LG/J) mouse, have been reported to regenerate multiple tissue types without fibrosis (Clark et al., 1998; Heydemann, 2012). The most well-studied example of MRL regeneration is that of ear punch wounds: while through-and-through ear wounds remain open and fail to regenerate the excised tissue in most mouse strains, MRL mice fully heal these wounds via initial formation of a blastema-like structure and subsequent regeneration of key tissue types including cartilage and hair-bearing skin (Clark et al., 1998). However, the molecular mechanisms underlying enhanced wound healing in the MRL ear remain poorly understood. To date, nine quantitative trait locus (QTL) mapping studies have been performed to identify associations between genomic regions and the ear closure phenotype (Blankenhorn et al., 2003, 2009; Cheverud et al., 2012, 2014; Heber-Katz et al., 2004; Masinde et al., 2001; McBrearty et al., 1998; Yu et al., 2005, 2007), but these studies so far have failed to identify specific genes or pathways driving regenerative healing, with ear closure-associated QTL spanning dozens or hundreds of individual genes.

Identification of tissue- and behavior-specific *cis*-regulatory divergence, through analyses of allele-specific gene expression in hybrids, has previously revealed genes and pathways underlying complex traits (Combs et al., 2018; Hu et al., 2022; York et al., 2018). MRL regeneration is wound site-specific – while ear wounds regenerate, dorsal wounds form fibrotic scars similar to other mouse strains (Colwell et al., 2006) – providing the opportunity to apply a similar approach to elucidate genes driving MRL ear regeneration. Here, we capitalize on mouse strain- and site-specific differences in healing to identify divergence in *cis*-regulation of gene expression associated with MRL ear regeneration through allele-specific expression (ASE) analysis. Collectively, our results highlight the power of this approach in dissecting complex phenotypes in mammals and implicate the complement pathway as a possible therapeutic target for improving wound healing and reducing scarring.

## Results

### Tissue- and strain- specific differences in wound healing

We first sought to robustly establish differences in healing phenotypes between two mouse strains: MRL, which regenerate ear wounds but heal dorsal wounds via scarring (Beare et al., 2006; Clark et al., 1998; Colwell et al., 2006); and CAST/EiJ (CAST), which do not possess any known strain-specific regenerative ability (Heber-Katz et al., 2004; Yu et al., 2005). Ear wounds were generated using a 2 mm punch tool to create a through-and-through wound in the center of the pinnae. Full-thickness dorsal excisional wounds were created via a previously published protocol (Mascharak et al., 2021a); in this model, silicone splints are applied around wounds to prevent the rapid contraction that typically occurs in mice and instead yield healing via granulation and re-epithelialization with human-like kinetics. Consistent with previous work (Clark et al., 1998; Yu et al., 2005), MRL ear wounds had largely closed by 3-4 weeks after wounding, with regeneration of normal-appearing skin grossly and cartilage histologically, while CAST ear wounds failed to close to an appreciable extent and instead formed scar tissue over the exposed wound edge (**Fig. 1A-B**). In contrast, dorsal wounds healed at a comparable rate in the two strains, with re-epithelialization complete by postoperative day (POD) 14 (**Fig. 1C-D**). Both MRL and CAST dorsal wounds healed by forming fibrotic scars, which grossly and histologically appeared as “bare areas” devoid of dermal appendages (e.g., hair follicles) and with dense connective tissue (**Fig. 1C**). Analysis of wound ECM ultrastructure using a previously published image analysis pipeline (Mascharak et al., 2022) confirmed that dorsal wounds in both CAST and MRL healed with ECM architecture that was quantitatively distinct from that of unwounded skin (consistent with fibrotic scar ECM); while CAST ear wounds also had a distinct ECM pattern, MRL ear wounds had ECM features largely overlapping with those of unwounded ear tissue, suggesting regeneration at the tissue ultrastructural level (**Fig. S1**).

**Figure 1:**
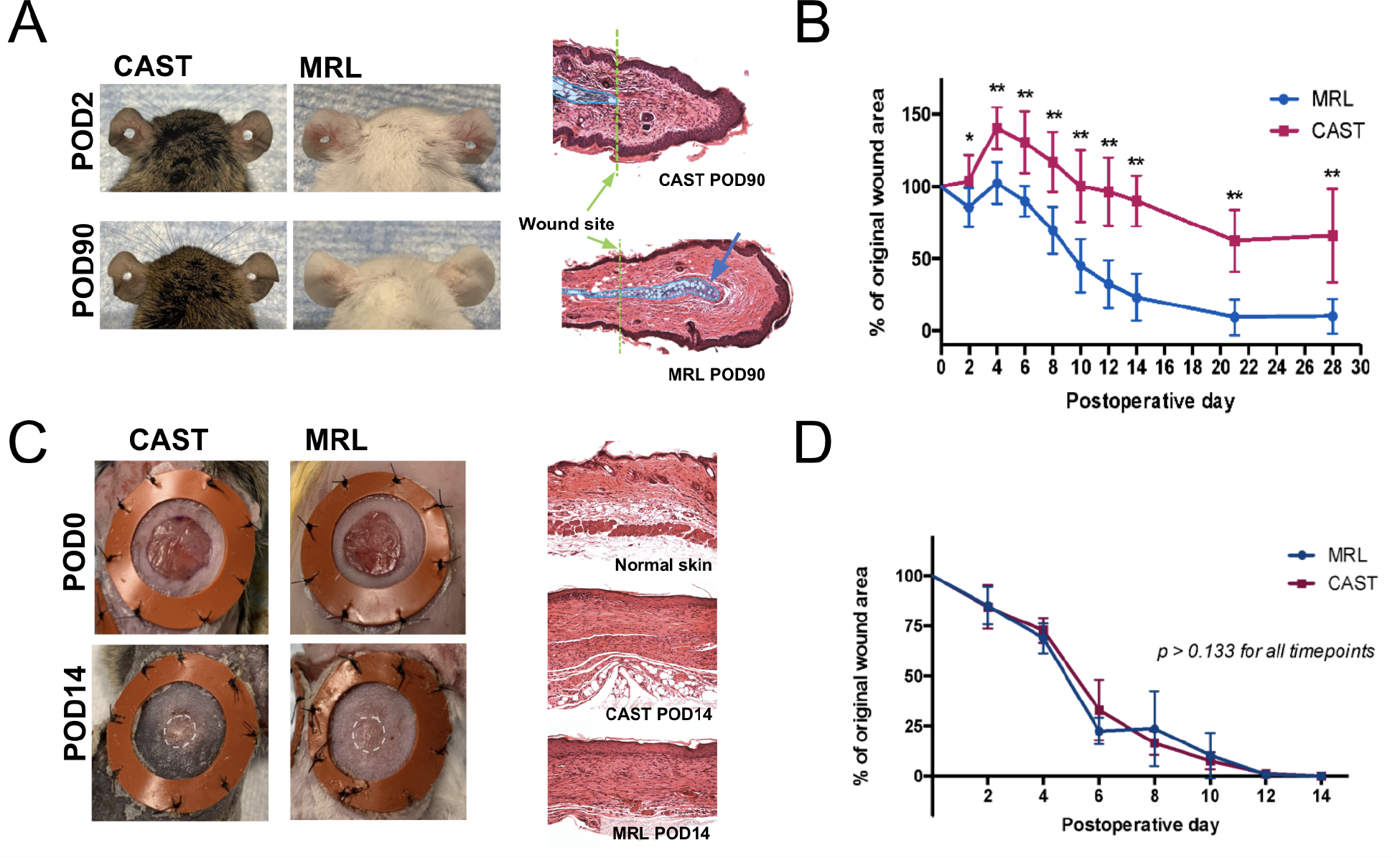
MRL ear wounds uniquely heal in an accelerated and regenerative fashion. A. Gross photographs (first two columns) at postoperative day (POD) 2 and 90 of CAST (“normal healer”) and MRL (“super healer”) mice. Hematoxylin and eosin (H&E) histology (third column) of ear wounds at POD 90. Green dotted lines indicate border of wound site; blue overlay indicates cartilage; blue arrow highlights regenerating cartilage in MRL ear wounds. **B.** Wound curves for CAST and MRL ear wounds showing closure over time. **C.** Gross photographs (first two columns) of splinted excisional dorsal wounds in CAST and MRL mice; white dotted lines indicate fibrotic scar “bare area.” H&E staining (third column) of POD 14 wounds and unwounded skin. **D.** Wound curves for CAST and MRL dorsal wounds reflecting rate of re-epithelialization over time***. B, D***. \**P* < 0.05, \*\**P* < 0.01 (Student’s t-test).

### Extensive *cis*-regulatory divergence during wound healing between regenerative and non-regenerative mouse strains

Having confirmed that enhanced/regenerative healing is specific to both the strain (MRL) and anatomic site (ear), we sought to leverage this unique phenotypic pattern to identify genes responsible for regeneration versus fibrosis. As wound repair involves a series of transcriptional cascades triggered by injury (Schäfer and Werner, 2007), we hypothesized that MRL ear closure may be driven by wound site-specific *cis*-regulatory activity. To identify *cis*-regulatory variation associated with wound healing, we crossed MRL with CAST mice to generate CAST x MRL F1 hybrids. In these first-generation hybrid offspring, alleles from both parents are present in the same cellular environment (i.e., are subject to the same *trans*-regulatory influences); so, differences in expression between the two alleles can only be the result of *cis*-regulatory differences (Babak et al., 2015; Cowles et al., 2002; Wittkopp et al., 2004). Thus, assaying allele-specific gene expression in F1 hybrid wounds allows for identification of injury-relevant *cis*-regulatory differences between these two mouse strains. Further, comparing allele-specific expression between wounds in the ear – which exhibits regenerative healing specifically in MRL mice – and the dorsum – which heals with a scar in both MRL and CAST – may allow us to pinpoint causal *cis*-regulatory differences driving the unique MRL ear regeneration phenotype.

In order to assess allele-specific gene expression across wound contexts, we performed RNA-seq of key cell populations associated with wound healing. Adult F1 female mice were subjected to dorsal splinted excisional and ear punch wounding. On POD 7, all wounds were harvested. Ear and dorsal wound tissue were separately digested and subjected to fluorescence-activated cell sorting (FACS) to isolate three cell populations: immune cells (CD45^+^); endothelial cells (CD31^+^); and fibroblasts (Lin^-^, per published sorting strategy(Leavitt et al., 2017); see Methods for details). Due to cell number limitations, wounds from three individual mice were pooled per biological replicate. Cell samples were then subjected to bulk RNA-sequencing (**Fig. 2A**). At least three biological replicates were sequenced and analyzed for each cell type.

**Figure 2:**
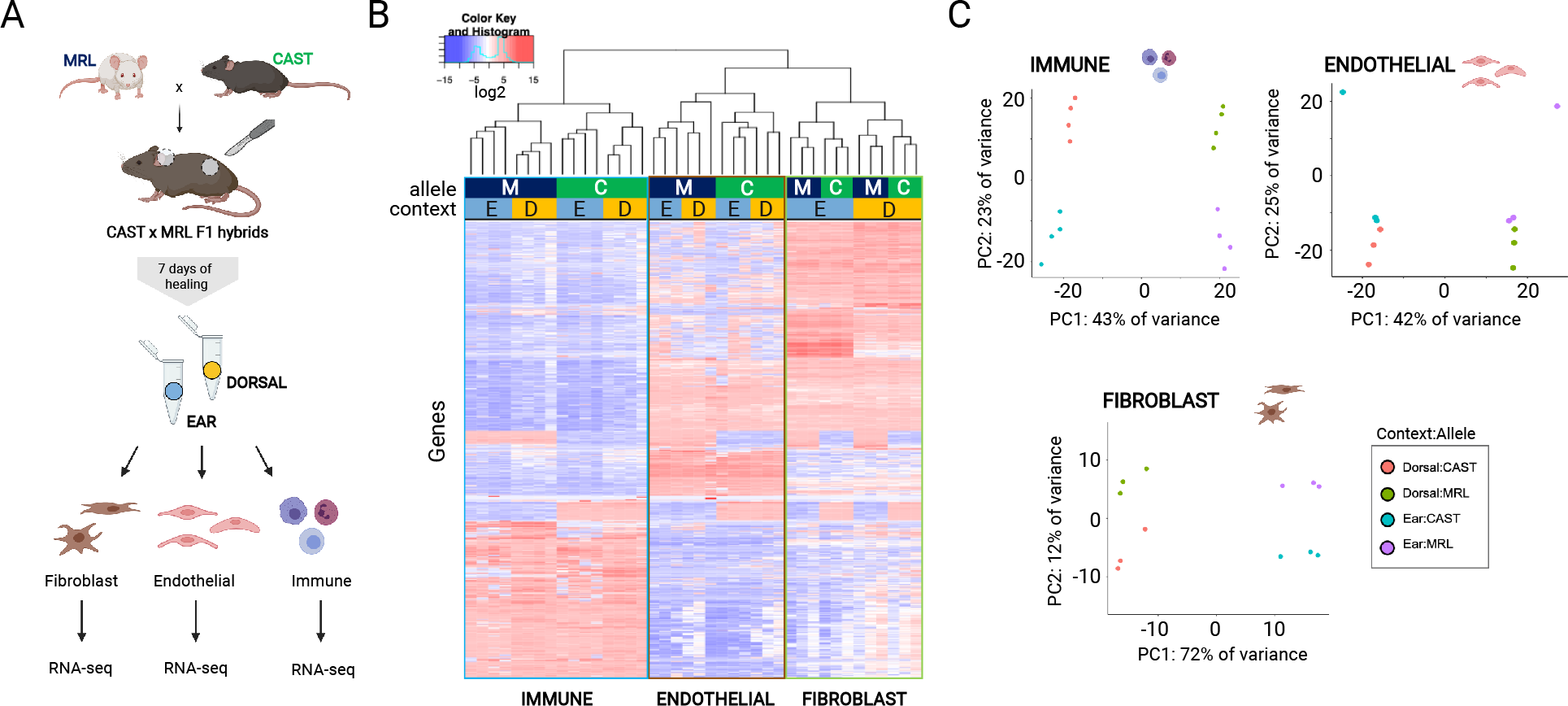
RNA-seq of key wound cell types from CAST x MRL hybrid mice cluster by wound type and allele. A. Sampling scheme for RNA-seq libraries. MRL and CAST were crossed to produce F1 hybrids for allele-specific expression analysis. Each adult F1 mouse underwent both dorsal excisional and ear punch wounding. On POD 7, wound tissue was harvested, cell populations were isolated via fluorescence-activated cell sorting (FACS), and RNA was extracted for bulk RNA-seq. **B.** Heatmap of the most variable genes (1,000) following regularized log2 transformation of allele-specific read counts. Hierarchical clustering groups samples by cell population (immune, endothelial, fibroblast), allele (MRL [‘M’] vs. CAST [‘C’] and wound site (ear [‘E’] vs. dorsal [‘D’]). **C**. Principal component analysis of allele-specific read counts. Allele-specific samples separated into distinct clusters by wound site (ear vs. dorsal) and allele (MRL vs. CAST) for each cell population.

Across all tissue samples, we obtained a total of ∼3.6 billion reads (**Table S1**). To enable the allele-specific assignment of RNA-seq reads, we performed whole-genome re-sequencing of MRL (∼25X depth of coverage; **Fig. S2**). Genetic variants differing between MRL and CAST strains were used to preferentially assign RNA-seq reads to either the MRL or the CAST allele (**Table S1**). After filtering for genes with low coverage in either genotype or wound context, we were able to analyze >10,000 genes in each cell population (**Table S2**). Hierarchical clustering and principal component (PC) analysis of allele-specific expression data grouped samples strongly by cell type (fibroblasts, immune, endothelial) (**Fig 2B**, **Fig. S3**). PC analysis of individual cell populations clearly separated samples by wound site (dorsal vs. ear: PC1 for fibroblasts, 71% of variance explained; PC2 for immune and endothelial; 23% and 25% of variance explained, respectively) and allele (CAST vs. MRL: PC2 for fibroblasts, 14% of variance explained, PC1 for immune and endothelial, 43% and 42% of variance explained respectively; **Fig. 2C**).

We next sought to identify genes with expression patterns reflecting MRL ear-specific *cis*- regulatory divergence, which could reflect functional involvement of these genes in driving regeneration rather than fibrosis for wounds in this tissue (**Fig. 3A**). Across all cell types, a greater number of genes exhibited significant allele-specific expression (ASE; FDR < 0.05 for MRL vs. CAST allelic expression with DESeq2) in ear wounds (5,121 genes; 32.7%) compared to dorsal wounds (2,655 genes; 17%), consistent with phenotypic divergence restricted to the ear (**Fig. 3B, Table S3**). This was most apparent in immune cells, where over four times as many genes had evidence of ASE in ear compared to dorsal wounds. Differences in expression between CAST and MRL alleles (i.e., |log2 fold change|) were also larger on average in ear wounds in each cell population (**Fig. S4**; Wilcoxon rank sum test, all ear vs. dorsal pairwise comparisons *p* < 2.2 x 10^-16^).

**Figure 3:**
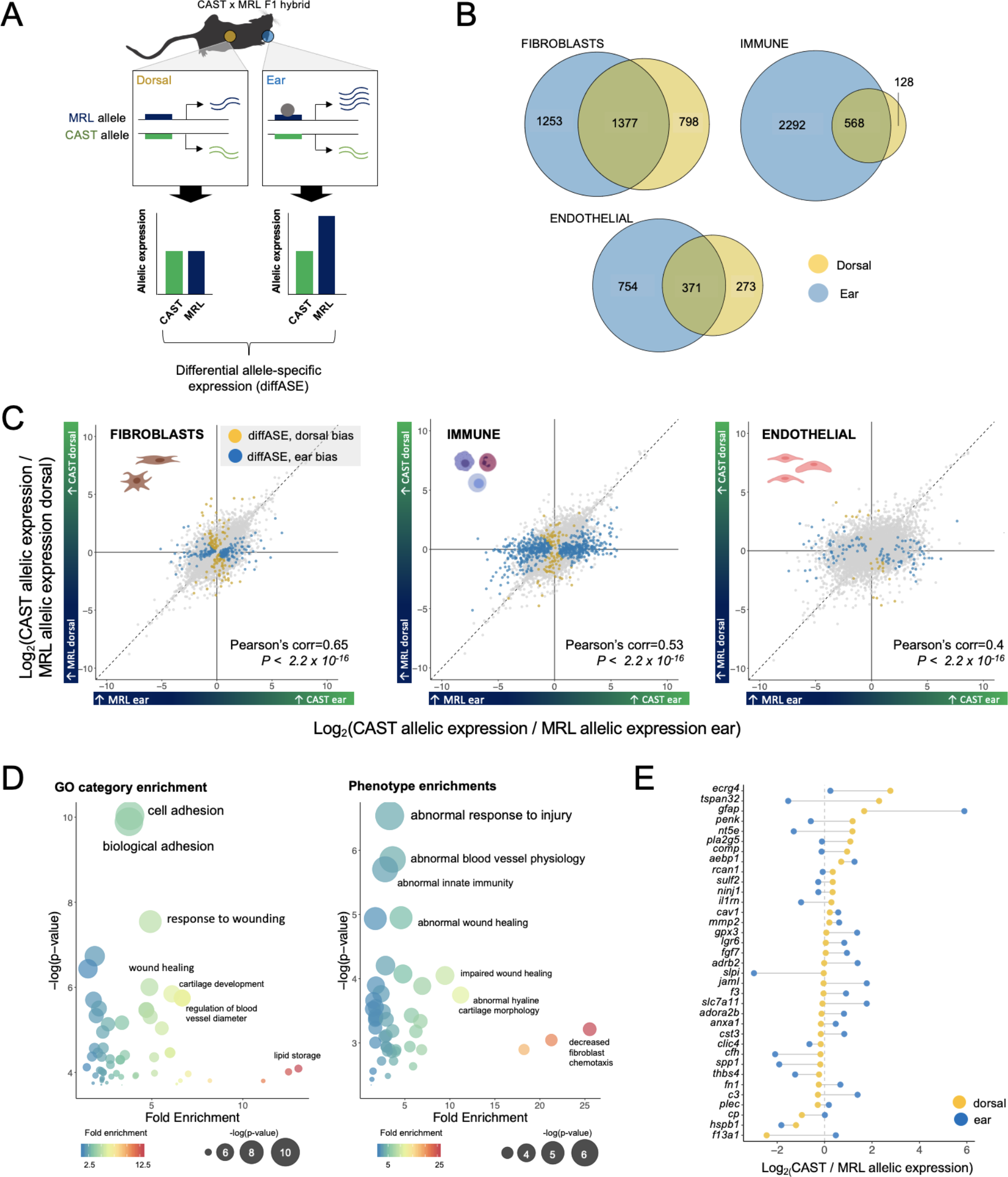
Analysis of differential allele-specific expression (diffASE) reveals *cis*-regulatory divergence unique to MRL ear wounds. A. Schematic example of diffASE between MRL and CAST in ear and dorsal wounds. Blue and green solid boxes represent gene regulatory regions affecting transcription of the MRL or CAST allele, respectively, of a given gene; transcription levels from each allele are represented by blue and green wavy lines. In the context of a dorsal wound (where MRL and CAST phenotypes are similar), expression is the same from the MRL vs. CAST allele. In contrast, in ear wounds, the presence of a context-specific (i.e., wound-related) transcription factor (TF; grey circle) reveals ASE through differences in the sequence of the MRL vs. CAST regulatory elements (which respond differentially to the TF). Overall, this results in a pattern of diffASE, where allele-specific expression is unique to ear wounds (exemplified in bottom panel bar graphs). **B.** Venn diagrams showing number of genes with ASE in ear wounds (blue region), dorsal wounds (yellow region), or both (overlapping region) in each analyzed wound cell type. **C.** Scatterplots for each cell type comparing distribution of allelic ratios between dorsal and ear wounds. Colored points represent genes with diffASE (gold points are genes with a larger difference between CAST and MRL alleles in the ear; blue points are genes with a larger difference between CAST and MRL alleles in the dorsum). **D.** Gene set enrichment analysis for genes with evidence of diffASE in fibroblasts, which are highly enriched for gene ontology (GO) categories (left) and mutant phenotypes (right) related to wound healing and injury responses. Such enrichment patterns were unique to fibroblasts (the end cellular mediators of scarring/fibrosis) and not seen in endothelial or immune cells. **E.** Specific genes associated with mutant phenotypes or GO terms related to responses to injury and wound healing with diffASE in fibroblasts. Yellow circles represent fold changes between alleles in the dorsum; blue circles represent fold changes in the ear.

Considering both ear and dorsal wounds, approximately 23% of genes demonstrated ASE in more than one cell population (**Fig. S5**). Within each cell population, while a substantial proportion of genes showed ASE in both wound contexts, many exhibited ASE unique to either ear or dorsal wounds (**Fig. 3B**), suggesting the existence of both general and tissue-specific regulatory divergence between CAST and MRL during wound repair. For genes with ASE in both dorsal and ear wounds, the vast majority maintained the same directionality of allelic expression across wound sites (i.e., same allele up-/down-regulated in both ear and dorsal wounds; **Table S4**). Further, we found that allelic ratios were correlated between wound sites (i.e., log2(CAST ear/MRL ear) vs. log2(CAST dorsal/MRL dorsal); Pearson’s correlation, all comparisons *p* < 2.2 x 10^-16^; **Fig. 3C**). Taken together, our findings of ASE were consistent with greater context-dependent regulatory divergence in ear wounds relative to dorsal wounds.

### Wound context- and strain- specific *cis*-regulatory activity identifies genes involved in wound healing and injury response pathways

As MRL mice demonstrate a regenerative phenotype unique to the ear wound context and not seen in dorsal wounds (Colwell et al., 2006) (**Fig. 1**), we reasoned that the subset of genes with differential allele-specific expression (“diffASE”) between ear and dorsal wounds could include genes driving the regenerative healing phenotype specifically in MRL ear wounds. Across different wound settings, context-specific ASE may reflect wound site-specific activity of genes controlled by *cis*-regulatory elements with sequence differences between MRL and non-regenerating (e.g., CAST) mice (**Fig. 3A**). Comparing genes’ ASE measurements in ear and dorsal wounds, we identified 432 genes in immune cells, 91 in endothelial cells, and 235 in fibroblasts with diffASE between wound healing contexts (DESeq2 Wald-test, [CAST/MRL ear counts] vs. [CAST/MRL dorsal counts], FDR < 0.05; see Methods; **Fig. 3C, Table S3**). The majority of genes with diffASE were unique to a single cell population (732/745 genes total).

Examining genes with diffASE in each cell population, we found that those in fibroblasts were uniquely enriched for known mutant phenotypes and gene ontology (GO) terms associated with injury and wound repair (**Fig. 3D**). In contrast, wound healing-related terms were not enriched for either immune or endothelial cell diffASE genes, suggesting that *cis*-regulatory divergence in fibroblasts may play a particularly important role in driving divergent wound healing phenotypes in the MRL ear versus dorsum. In fibroblasts, genes with diffASE were most highly enriched for the mouse mutant phenotype “abnormal response to injury” (MP:0005164, FDR-adjusted *p*-value = 1.85 x 10^-5^) and were also enriched for the phenotypes “abnormal wound healing” (MP:0005164, FDR = 0.00086) and “abnormal blood vessel physiology” (MP:0000249, FDR = 0.00023). Additionally, the GO terms “response to wounding” (GO:0009611, FDR *=* 1.37 x 10^-4^) and “wound healing” (GO:0042060, FDR *=* 2.37 x 10^-3^) showed greater than four-fold enrichment compared to a background set. Fibroblast diffASE genes were also enriched for GO and Reactome Pathway terms related to processes involved in scarring and regeneration. These included cell adhesion (GO:0007155, FDR = 1.34 x 10^-6^) and integrin cell surface interactions (R-MMU- 216083, FDR = 1.95 x 10^-2^), which are implicated in activated mechanotransduction and pro-fibrotic changes in fibroblasts (Agarwal, 2014; Mascharak et al., 2021b); and extracellular matrix organization (GO:0030198, FDR = 3.50 x 10^-3^; and R-MMU-1474244, FDR = 9.52 x 10^-3^), a critical determinant of scarring versus regenerative wound properties, among others (**Fig. 3D**; full list of enriched terms in **File S1**).

Further, we identified several genes associated with wound repair phenotypes or known pathways with large differences in allelic ratio between ear and dorsal wound fibroblasts (**Fig. 3E, Table S5, Table S6**). Some of these genes with MRL-specific upregulation in ear wounds had known functions consistent with promoting wound healing and/or decreasing scarring. For instance, *Slpi* (Secretory leukocyte protease inhibitor; ear wounds: log2(CAST/MRL) = -2.98, *q* = 5.74 x 10^-6^; dorsal wounds: log2(CAST/MRL) = -0.036, *q* = 0.58) has important functions in wound healing, in part via regulating transforming growth factor-beta (TGF-β) activity, and *Slpi*-null mice exhibit delayed wound repair and increased inflammation (Ashcroft et al., 2000; Zhu et al., 2002). *Spp1* (Secreted phosphoprotein 1, which encodes the protein osteopontin; ear: log2(CAST/MRL) = -1.91 FDR = 1.083 x 10^-36^; dorsal: log2(CAST/MRL) = -0.18, FDR = 0.57) has been implicated in resolution of inflammation as well as matrix remodeling following skin injury (Bevan et al., 2020; Liaw et al., 1998), the latter being especially critical in determining scarring versus regenerative healing outcomes (Mascharak et al., 2022). Additionally, osteopontin knockout mice have impaired wound closure (Wang et al., 2017). *Thbs4* (Thrombospondin 4, an ECM protein; ear: log2(CAST/MRL) = -1.24, FDR = 6.55 x 10^-38^; dorsal: log2(CAST/MRL) = -0.23, FDR = 0.048) has previously been shown to promote wound healing by stimulating fibroblast migration and keratinocyte proliferation (Klaas et al., 2021) and is reported to promote angiogenesis and reduce fibrosis (Stenina-Adognravi and Plow, 2019), with mouse *Thbs4* knockout associated with damaging cardiac inflammation and fibrosis (Frolova et al., 2012).

We also identified several genes with reported functions that promote fibrosis and/or impair injury repair, which exhibited upregulation from the CAST allele in ear wounds. For instance, *Jaml* (Junctional adhesion molecule-like; ear: log2(CAST/MRL) = 1.78, *q* = 4.24 x 10^-7^; dorsal: log2(CAST/MRL) = -0.04, *q* = 0.87) has been associated with inflammation and impaired injury repair in the intestine (Parkos et al., 2013; Weber et al., 2014). *Adora2b* (Adenosine A2b receptor; ear: log2(CAST/MRL) = 0.82, FDR = 0.00057; dorsal: log2(CAST/MRL) = -0.12, FDR = 0.71) inhibition has been associated with reduced dermal fibrosis (Karmouty-Quintana et al., 2018), consistent with a pro-scarring role for this gene. Collectively, many genes exhibiting diffASE preferentially in ear wounds had known functions consistent with the phenotypes observed in CAST versus MRL ear wounds (i.e., pro-fibrotic genes enriched from the CAST allele; pro-regenerative genes and genes promoting efficient wound repair enriched from the MRL allele).

### Overlap with healing quantitative trait loci identifies candidate genes for regeneration

Next, we sought to integrate our results with prior functional studies of MRL ear regeneration. Specifically, having identified genes with wound site-specific *cis*-regulatory differences, we next asked whether those genes were located within previously mapped genomic intervals associated with enhanced ear punch closure. We capitalized on a recent QTL fine-mapping study for ear wound closure in LG/J (LG), the MRL progenitor line (LG x SM, F32 generation) (Cheverud et al., 2014). LG shares ∼75% of its genome with MRL and exhibits similar regenerative healing of ear punch wounds (Cheverud et al., 2012; Li et al., 2001). Consequently, overlap between these studies will be restricted to causal loci for regenerative healing that are shared between lines. To test whether our diffASE gene sets were enriched in genomic regions driving ear wound closure, we compared the LOD scores of genetic markers closest to genes with diffASE (**Fig. 4A**) against those of randomly permuted gene sets of the same size. While the presence of a significant LOD score proximal to a single gene does not necessarily implicate that gene, a shift towards a higher average LOD score for a group of genes suggests that this set of genes is collectively more likely to be associated with differences in wound phenotypes (i.e., regenerative versus non-regenerative). Our analysis revealed that genes with diffASE in fibroblasts had significantly higher average LOD scores compared to random sets (20,000 permutations, diffASE genes FDR < 0.05, *p* = 0.0094; diffASE FDR < 0.1, *p* = 0.026; **Fig. 4B**). In contrast, genes with diffASE in immune or endothelial cells did not exhibit higher LOD scores on average (*p* > 0.05 for each comparison; 20,000 permutations). Further, genes with ASE in *both* ear and dorsal wound fibroblasts were not associated with higher LOD scores (*p* > 0.05), suggesting that diffASE was uniquely useful in pinpointing causal wound healing genes.

**Figure 4:**
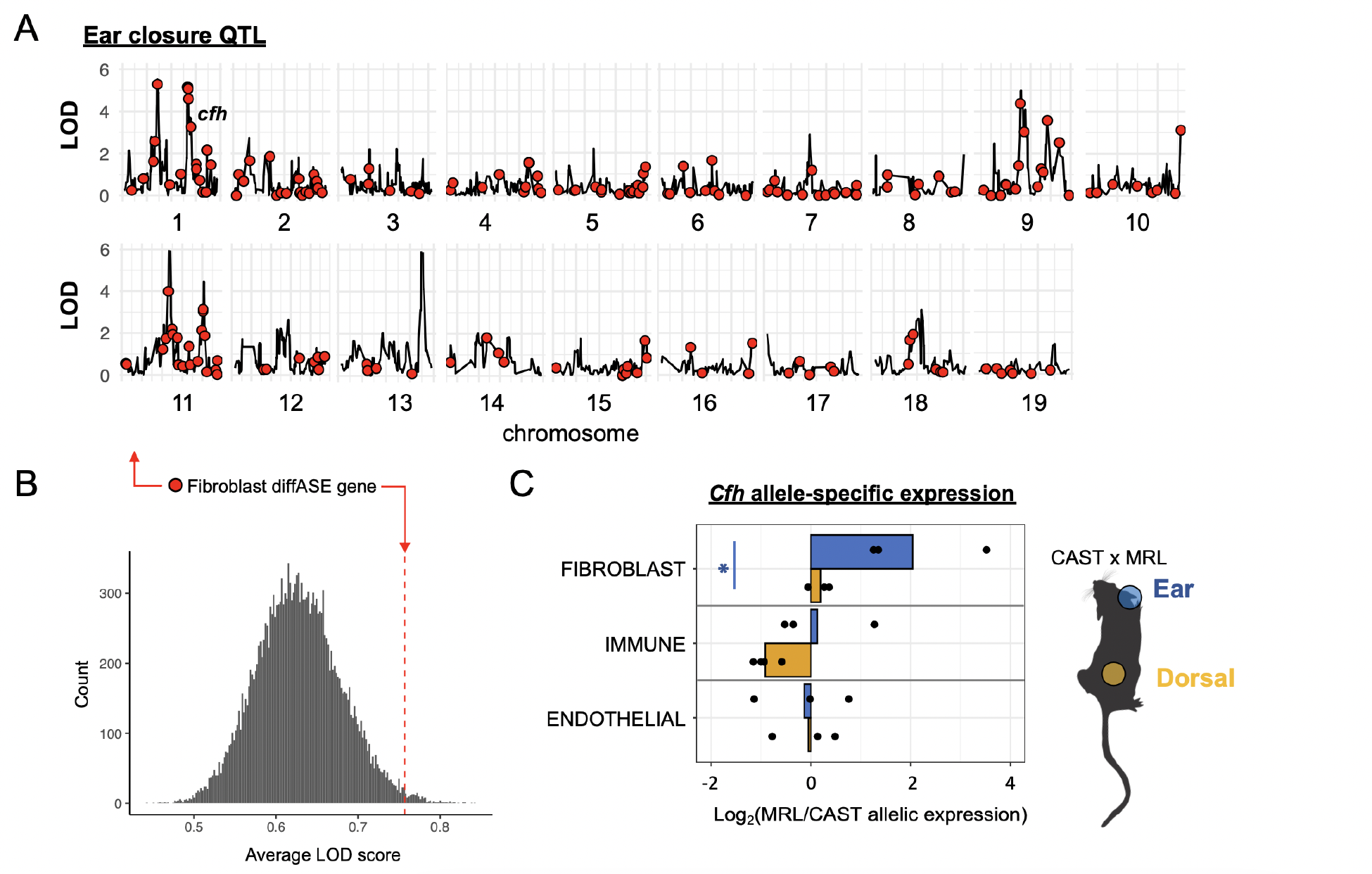
Integration of diffASE with QTL fine-mapping study identifies *Cfh* as a candidate gene for driving the MRL regenerative healing phenotype. A. LOD scores vs. chromosome position for ear hole closure from Cheverud et al. 2014. Red circles indicate the positions of genetic markers closest to genes identified as having diffASE in fibroblasts. **B**. Distribution of mean LOD scores of permuted gene sets (20,000 permutations). Red line indicates the mean LOD score of genetic markers closest to the fibroblast diffASE gene set. C. *Cfh*, which is associated with the gene ontology term for wound healing (GO:0042060) and falls within a fine-mapped region for ear closure, shows ear wound-specific ASE specific to fibroblasts in CAST x MRL hybrids. In fibroblasts, we see significant upregulation of the MRL allele relative to the CAST allele in ear wounds, in contrast to dorsal wounds where the expression of these alleles are similar. *diffASE *q* < 0.05.

Next, to identify specific candidates for driving regenerative wound healing, we searched for genes with diffASE located within support intervals of significant wound closure QTL. Across cell types, we identified a total of 27 genes with diffASE overlapping these QTL intervals (diffASE FDR < 0.05; fibroblasts, 9 genes; endothelial cells, 6 genes; immune cells, 12 genes; diffASE FDR < 0.1, 40 genes; **Fig. S6, File S1**). Genes with diffASE in fine-mapped intervals were enriched for the GO term “wound healing” (GO:0042060; Fisher’s exact test, *p* = 0.0032, 9.45-fold enrichment). Several genes within these intervals could be promising candidates based on their known functions or mutant phenotypes (see **File S1**); specifically, three of the genes with diffASE in fibroblasts – *Fn1, Lgr6,* and *Cfh* – were associated with ontology terms for wound healing and regeneration. Of these genes, *Cfh* (Complement factor H) had the greatest difference in allelic ratios between ear and dorsal wounds, with expression from the MRL allele over four times that of the CAST allele on average in ear wounds, but no significant difference between alleles in dorsal wounds (**Fig. 4C**). The complement cascade is a part of the innate immune system that is involved in clearing microbes, immune complexes, and damaged self cells and is activated in response to tissue injury (Cazander et al., 2012; Józsi and Zipfel, 2008). In addition to *Cfh*, ear-specific ASE in wound fibroblasts was also observed in two other genes encoding complement components, *C3* and *C1qb*, both of which have been directly implicated in wound repair (Bossi et al., 2014; Hayuningtyas et al., 2021; Rafail et al., 2015). While the complement system plays a vital role in injury repair – for example, *C1qb* is important for angiogenesis (Bossi et al., 2014) – inappropriate or prolonged complement activation is also known to perpetuate damaging inflammation and cause cell death (Cazander et al., 2012). However, prior studies conflict on whether complement activation or inhibition may promote wound healing (Rafail et al., 2015; Sinno et al., 2013), and the effects of complement modulation on scarring have not been investigated.

### Ectopic application of CFH reduces scarring and drives partial regeneration after wounding

Complement factor H is a central regulatory protein in this pathway that inhibits complement activation (Józsi and Zipfel, 2008) and which has previously been shown to prevent inflammation and fibrosis in the mouse kidney (Alexander et al., 2005). Given these known functions, strong ear wound-specific ASE of *Cfh*, and its presence within a QTL interval for ear closure, we hypothesized that *Cfh* could be a driver of wound regeneration. We first sought to verify that our gene expression findings corresponded to differences at the protein level. First, we cultured fibroblasts from the ear and dorsum of both CAST and MRL mice, then performed immunostaining for CFH (**Fig. 5A**). CFH protein expression was absent in both ear and dorsal CAST fibroblasts, and was significantly greater in MRL ear than MRL dorsal fibroblasts (**Fig. 5B**), consistent with our finding of MRL ear-specific upregulation from RNA-seq. We next performed CFH staining on sections from POD 14 ear and dorsal wounds from CAST and MRL (**Fig. 5C**), which revealed that CFH protein expression was markedly upregulated in MRL ear wounds compared to all other conditions (**Fig. 5D**).

**Figure 5:**
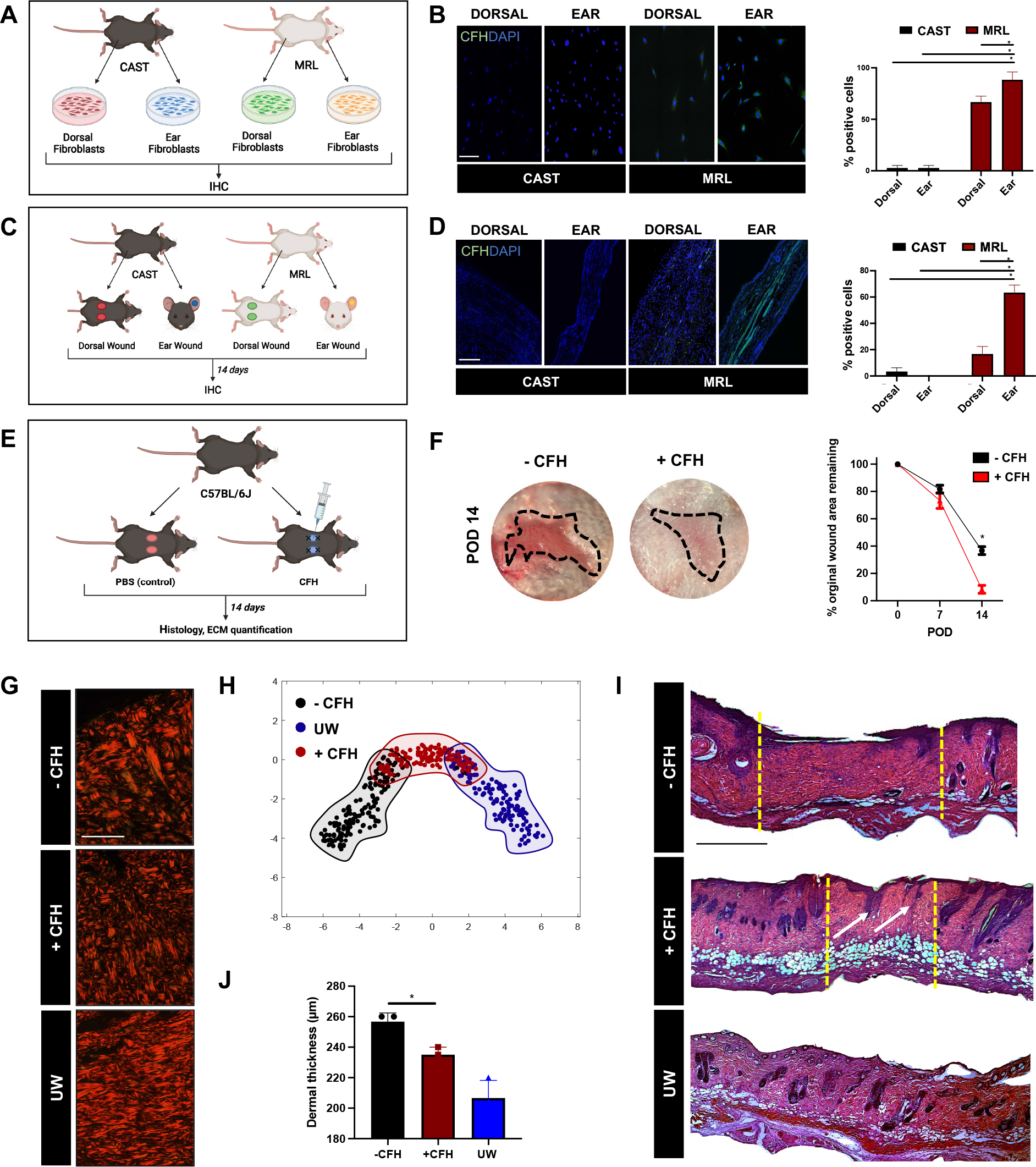
CFH treatment leads to partial regeneration and enhanced healing of dorsal wounds in wildtype mice. A. Schematic of MRL and CAST dermal fibroblast culture from dorsal and ear skin. **B.** Left, fluorescent histology of cultured MRL and CAST dorsal and ear fibroblasts with immunohistochemical (IHC) staining for complement factor H (CFH) and DAPI nuclear counterstain. Right, quantification of CFH expression across *in vitro* conditions. **C.** Schematic of MRL and CAST dorsal and ear wounding for histology. **D.** Fluorescent histology (left) and quantification (right) of IHC staining of wounds for CFH (with DAPI nuclear counterstain). **E.** Schematic of wildtype mouse dorsal splinted wounding with local wound treatment with either recombinant CFH protein or phosphate-buffered saline (PBS; vehicle control) (see Methods for full details and dosing). **F.** Left, gross photographs of control (-CFH) and CFH-treated (+CFH) wounds; black dotted outline indicates healed wound region. Right, wound curve reflecting rate of re-epithelialization of -CFH vs. +CFH wounds over time. **G.** Picrosirius red connective tissue histology of -CFH and +CFH wounds and unwounded skin (UW). **H.** T-distributed stochastic neighbor embedding (t-SNE) plot of quantified extracellular matrix (ECM) ultrastructural parameters, based on picrosirius red histology of unwounded skin and POD 14 wounds (**G**), showing overall similarities/differences in ECM ultrastructure between conditions. Each dot represents quantified parameters from one histologic image. **I.** Hematoxylin and eosin (H&E) histology of POD 14 wounds and skin. Yellow dotted lines denote borders of healed wounds; white arrows indicate putative regenerating dermal appendages (hair follicles or glands) in +CFH wounds. **J.** Dermal thickness quantified from histology of wounds and skin. ***B, C, F, J.*** \**P* < 0.05 (Student’s t-test).

Given that CFH expression was strongly associated with regenerating conditions (MRL ear wounds), we next evaluated whether modulating CFH signaling could drive wound regeneration and reduce scarring. We treated dorsal wounds in another scarring wildtype mouse strain (C57BL/6J) with recombinant CFH protein or vehicle control (phosphate-buffered saline [PBS]), then evaluated wound outcomes (**Fig. 5E**; see Methods for full details and dosing). Grossly, CFH-treated wounds had reduced scarring and more robust re-epithelialization by POD 14 compared to control wounds (**Fig. 5F**). Quantification of ECM ultrastructural parameters from picrosirius red histology (**Fig. 5G**) showed that CFH treatment yielded ECM intermediate between that of control scars and unwounded skin (**Fig. 5H**), indicating partial regeneration. On hematoxylin and eosin (H&E) histology, CFH-treated wounds had more complete re-epithelialization (confirming gross observations), significantly reduced dermal thickness (consistent with reduced scarring), and structures morphologically resembling early invaginating neogenic hair follicles, in contrast with control scars which remained “bare areas” devoid of any dermal appendages (**Fig. 5I-J**). Finally, we also found that the reduction in scar thickness with CFH treatment was dose-dependent, with more significant scar prevention observed at higher doses (**Fig. S7**; see Methods for details). Collectively, these findings were consistent with CFH being sufficient to drive partial wound regeneration and significantly reduce scarring in mouse dorsal wounds, and suggest that this gene may play a similar, pro-regenerative role in MRL ear wounds.

## Discussion

Healing via fibrosis, rather than through regeneration, is a major cause of morbidity and an immense burden for healthcare systems worldwide, with over $20 billion spent annually on the treatment and management of scars in the United States alone (Block et al., 2015). Instances of regeneration in nature, such as the striking example of regenerative mammalian wound repair that occurs in the ears of MRL mice, may provide valuable insights for therapeutically promoting regeneration and preventing fibrosis.

While QTL mapping has previously been used to identify genomic intervals associated with regenerative healing in the MRL strain, identifying specific candidates for functional followup has proven challenging, in part due to the large size of intervals identified. Our approach, using allele-specific gene expression to probe for site-specific *cis*-regulatory divergence, offers the advantage of not only interrogating potential drivers of regeneration at the single-gene level, but also being substantially less resource-intensive (e.g., <20 mice used, compared to multiple hundreds in prior studies) and thus more accessible. The strength of our methodology is supported by multiple interesting findings. Our approach found greater ASE in ear compared to dorsal wounds across all cell types studied, as well as an enrichment of wound repair pathways and genes associated with differential allele-specific expression in fibroblasts. This would not be expected if the genetic changes leading to MRL ear regeneration were entirely protein-coding, and instead suggests that the strain-specific phenotypic difference seen in ear wounds may reflect *cis*- regulatory divergence between CAST and MRL. Furthermore, our findings were supported by integrating our results with the QTL fine-mapping study by Cheverud et al. 2014. The overlap observed with this orthogonal method suggests that at least some of the *cis*-regulatory changes we identified underlie the MRL ear regeneration phenotype.

Of note, while our findings highlighted fibroblasts as the most likely drivers of MRL ear wound regeneration, our and other studies have implicated alterations in diverse cell types (e.g., reduced inflammation mediated by immune cells, more rapid re-epithelialization mediated by keratinocytes) in MRL ear regeneration. These could result from tissue-specific *cis*-regulatory differences directly affecting other cell types (such as immune or epithelial cells). However, they also likely result, at least in part, from cell-cell crosstalk mediated by fibroblasts. For instance, intimate fibroblast-keratinocyte crosstalk is critical for wound repair (Werner et al., 2007), and we have previously found that modulating pro-fibrotic fibroblast molecular processes can also induce changes in the overlying epidermis (Mascharak et al., 2022). Fibroblast-immune and immune-epithelial interactions have also been extensively reported in the context of injury repair (Leoni et al., 2015; Van Linthout et al., 2014). Thus, it is feasible that altered fibroblast phenotype in wounds could fundamentally drive many of the differences observed in regenerating MRL ear wounds; we found that fibroblast diffASE genes were enriched for pathways involved in modulating cartilage development and immune cell activity, further supporting this hypothesis.

Finally, our genomic findings are also supported by the results of our experiments with CFH, which was identified by diffASE analysis and subsequently shown via *in vivo* wounding experiments to promote regeneration and reduce scarring. In addition to providing important functional validation for our methodological approach, this finding could have important translational implications, as no targeted molecular therapies currently exist to prevent human scarring.

## Limitations of study

While the pro-regenerative effects of recombinant CFH treatment were substantial, they also fell short of complete regeneration. This is unsurprising, given that the molecular basis of wound regeneration in MRL mice is complex and known to involve many genomic regions (Heber-Katz et al., 2004; Masinde et al., 2001; McBrearty et al., 1998). Future studies may seek to determine whether other genes identified by our diffASE analysis also have similar pro-regenerative effects.

## Acknowledgments

We thank J. Cheverud for providing data from the wound closure QTL analysis. Funding for this work was provided by NIH R01-GM136659 (to M.T.L.), R01-GM097171 (to H.B.F.), U24- DE029463 (to M.T.L.); the Wu Tsai Human Performance Alliance, the Hagey Laboratory for Pediatric Regenerative Medicine, the Gunn Olivier Fund, the Scleroderma Research Foundation, and the Pitch and Catherine Johnson Fund (to M.T.L.); the Stanford Medical Scientist Training Program (to H.E.T.); and the Ruth L. Kirschstein National Research Service Award Individual Postdoctoral Fellowship (F32)(to K.L.M.).

## Author contributions

Conceptualization: HET, MJB, HBF, MTL Methodology: HET, KLM, HBF, MTL

Investigation: HET, KLM, MG, NJG, JBLP, AFS, MFD, SM, MJB

Visualization: HET, KLM, MG

Funding acquisition: HET, KLM, HBF, MTL Supervision: DCW, HBF, MTL

Writing – original draft: HET, KLM

Writing – review & editing: DCW, HBF, MTL

## Data availability

Illumina sequencing data from this study has been deposited to the NCBI Sequence Read Archive Raw (PRJNA839777). Read counts and gene lists for each cell type are available in **File S1** and on **Figshare** (https://doi.org/10.6084/m9.figshare.c.6025157).

## Materials availability

This study did not generate new unique reagents.

## Lead contact

Further information and requests for resources and reagents should be directed to the lead contact, Michael Longaker.

## Methods

### Mice

Mice were housed and maintained in sterile micro-insulators at the Stanford University Comparative Medicine Pavilion in accordance with Stanford University Administrative Panel on Laboratory Animal Care (APLAC) guidelines (APLAC-21308). Food and water were provided *ad libitum*. MRL/MpJ (MRL), CAST/EiJ (CAST), and C57BL/6J mice were obtained from The Jackson Laboratory (Bar Harbor, ME). Male MRL and female CAST mice were bred to produce CAST x MRL F1 hybrid (F1) offspring. Adult (postnatal day [P]60) female F1 mice were used for RNA-seq experiments, and both female and male P60 mice were used for all other experiments.

### Dorsal and ear wounding

Mice underwent ear punch wounding and dorsal splinted excisional wounding following established protocols without modification (Clark et al., 1998; Mascharak et al., 2021a). Briefly, mice were anesthetized using 1.5-2% isoflurane. All surgical tools were autoclaved prior to the procedure. For ear wounds, the skin was prepped using alcohol wipes. Punch wounds 2mm in diameter were created using a thumb-type metal ear punch (Fisherbrand). For mice wounded for histologic and wound curve analysis, one wound was created per ear, roughly in the middle of the pinna. For mice wounded for RNA-seq analysis, in order to obtain sufficient cells for analysis while minimizing the number of mice required, three wounds were created per ear, spaced at least 2mm apart. For dorsal wounds, hair was removed from the entire dorsum using an electric shaver followed by depilatory cream. The dorsal skin was then prepped using three sequential alternating swabs of betadine and 70% ethanol. Using sharp surgical scissors, two 6mm-diameter full-thickness excisional wounds were made per mouse, roughly at the level of the scapulae and 4mm lateral to midline. Wounds were stented open by affixing silicone rings (1cm internal diameter) around the wound using adhesive and eight simple interrupted sutures (6-0 nylon, Ethicon). Wounds were dressed using Tegaderm (3M) and dressings were changed every other day until harvest.

For dose response experiments with CFH-treated dorsal wounds (see Fig. S7), which were performed in CAST mice, recombinant mouse complement factor H protein (R&D Systems) was resuspended in phosphate-buffered saline (PBS) at a concentration of either 5 or 10 µg/mL, then 50 µL of CFH at these concentrations, or PBS (vehicle control), were injected locally into the wound base and surrounding dermis immediately following wounding (POD 0) and again at POD 7. For CFH scar prevention experiments (Fig. 5), which were done in C57BL/6J mice (due to poor anesthesia tolerance and excessive fighting leading to high morbidity/mortality with initial wounding experiments in CAST mice), recombinant CFH was resuspended at 30 µg/mL and 30 µL of CFH solution or PBS were injected on POD 0, 2, 5, and 7. For wound curve analysis, wounds were photographed every other day for the first two weeks of healing, then (for ear wounds) weekly for two additional weeks. For ear wounds, a circular stencil was placed over wounds prior to photographing in order to provide a consistently sized reference for measurements; for dorsal wounds, the silicone splints served as a size reference. Area of the stencil/splint and remaining wound area at each timepoint were measured in Photoshop (Adobe) and used to calculate remaining wound area as a percentage of original wound size (normalized to size of stencil/splint for each photograph).

#### Tissue histologic analyses

Tissue for histologic analysis was harvested and fixed by incubation in 10% neutral buffered formalin for 16-18 hours at 4 °C. Following fixation, tissue was processed for paraffin or OCT embedding by standard procedures. Briefly, for paraffin embedding, tissue underwent sequential dehydration (ethanol), clearing (xylene), and infiltration by paraffin wax. For OCT embedding, tissue was incubated in 30% sucrose/PBS for two weeks at 4 °C, OCT for 1 day at 4 °C, then embedded in OCT blocks by freezing in a dry ice/*tert*-butanol bath. All wounds were bisected and embedded cut-side-down. Tissue sections were cut using a microtome (paraffin) or cryostat (OCT) at 8 µm thickness. Hematoxylin and eosin (H&E) and picrosirius red staining (using Picro Sirius Red Stain Kit, Abcam) were performed on paraffin sections, using standard protocols without modification. Dermal thickness was measured from H&E histology images; Photoshop (Adobe) was used to measure the dermis (from the bottom of the epidermis to the top of the subcutaneous tissue), and a minimum of nine measurements (three measurements per image from three individual histology images/sections) were averaged per wound. Machine learning analysis of ECM ultrastructure was performed as previously described using Matlab (Mascharak et al., 2022). Briefly, picrosirius red histology images were normalized, color deconvoluted, noise reduced, then binarized. Binarized images were filtered to select for fiber-shaped objects and the fiber network was skeletonized. Finally, 294 parameters of the digitized map (including fiber length, width, persistence, alignment, etc.) were measured. Dimensionality reduction of quantified fiber network properties by t-distributed stochastic neighbor embedding (t-SNE) was used to plot parameters for each image. Comparisons between conditions were based on visual assessment of t-SNE clustering from calculated ECM parameters. Matlab scripts containing our fiber quantification pipeline are available at the following Github repository: https://github.com/shamikmascharak/Mascharak-et-al-ENF.

#### FACS isolation of wound cell populations

To harvest a consistent region for RNA-seq analysis, wounds were excised with a 1mm ring of tissue around each wound using a biopsy punch (4 mm punch for ear wounds; 8 mm punch for dorsal wounds). Wound tissue was incubated in ammonium thiocyanate (3.8% in Hank’s balanced salt solution [HBSS]) for 20 minutes at room temperature to dissociate the epidermis, then dermal tissue was separated from overlying epidermis and underlying cartilage (for ear wounds) under a surgical microscope. For each biological replicate, ear or dorsal wounds from three individual mice were pooled to obtain sufficient cell numbers for sequencing. Wounds were finely minced with sharp surgical scissors then enzymatically digested in collagenase type IV (1500 U/mL in Dulbecco’s Modified Eagle Medium [DMEM]) at 37 °C, with agitation at 150 rpm, for 1 hour. After 1 hour, digestion was quenched by addition of equal volume of DMEM with 10% heat-inactivated fetal bovine serum (FBS), filtered through 70 µm followed by 40 µm nylon filters, then pelleted (200 x *g*, 5 min, 4 °C). Cell pellets were resuspended in FACS buffer (PBS with 1% FBS and 1% penicillin-streptomycin) then stained with the following antibodies: PE anti-CD45 (BioLegend #103105); APC anti-CD31 (Invitrogen #17-0311-80); and eFluor 450-conjugated Lineage (Lin) antibodies anti-CD45 (ThermoFisher Scientific #48-0451-82), anti-Ter-119 (ThermoFisher Scientific #48-5921-82), anti-CD31 (BioLegend #303114), anti-Tie-2 (ThermoFisher Scientific #13-5987-82), anti-CD326 (ThermoFisher Scientific #48-5791-82), and anti-CD324 (ThermoFisher Scientific #13-3249-82), for isolation of fibroblasts via lineage depletion as per previously published protocol (Leavitt et al., 2017). DAPI (BioLegend; 1:1000) was added as a viability stain. Live (DAPI^-^) singlet cells were sorted to obtain immune cells (PE/CD45^+^), endothelial cells (APC/CD31^+^), and fibroblasts (PB/Lin^-^), which were sorted directly into lysis reagent (QIAzol, QIAGEN), then stored at -20 °C until RNA purification.

### Bulk RNA-sequencing of wound cell populations

RNA was purified from each cell sample using the miRNeasy Micro Kit (QIAGEN), then kept at -20 °C until sequencing. Samples were shipped on dry ice and library preparation and sequencing were performed by Admera Health (South Plainfield, NJ). Library preparation was with the SMART-Seq v4 Ultra Low Input Kit (Takara Bio) with PolyA Selection. Sequencing was done using Illumina HiSeq (2x150).

### Sequencing of MRL and variant calling

To identify variant calls for allele-specific expression, we generated whole-genome data for the MRL inbred line. MRL tail DNA was extracted and purified using the Invitrogen PureLink Genomic DNA Mini Kit. The library was prepared using the KAPA Hyper Prep kit. The MRL genome was sequenced to moderate coverage (average of 25x for sites with at least one read) on the Illumina HISeq X platform (2x150 reads; Fig. S2). Genomic reads were then mapped to the *M. m. domesticus* mm10 (GRCm38) reference genome using bowtie2 v2.3.4 (argument: --very-sensitive)(Langmead and Salzberg, 2012). CAST/EiJ sequence data was obtained from Wellcome Trust Mouse Genome Project (https://www.sanger.ac.uk/science/data/mouse-genomes-project)(Keane et al., 2011) in bam format, mapped to mm10. SNP calling was performed using the Genome Analysis Toolkit v4.1 (GATK)(McKenna et al., 2010). Duplicate reads were marked with the Picard tool MarkDuplicates. GATK HaplotypeCaller was used to call variants between CAST/EiJ, MRL, and the mouse reference genome (mm10). We filtered variants for low quality calls (SNPs: QD < 2.0 || FS > 60.0 || MQ < 40.0 || MQRankSum < -12.5 || ReadPosRankSum < - 8.0; Indels: QD < 2.0 || FS > 200.0 || ReadPosRankSum < -20.0), sites with a read depth of less than 5, and for any sites with heterozygous calls using GATK SelectVariants and bcftools (Danecek and McCarthy, 2017). This resulted in 19,472,153 SNPs, 1,171,415 of which were in exons. Variant calls where CAST and MRL differed from each other or both differed from the mm10 reference were then used to create alternative references for MRL and CAST for mapping. SNP calls were inserted in mm10 and indels were masked. The concatenated MRL-CAST genome was used to identify reads mapping uniquely to each parental genome in F1 hybrids.

#### Mapping and allele-specific assignment

Raw reads were trimmed for adapter contamination with the TrimGalore (v0.5) wrapper for Cutadapt v1.18 (Martin, 2011). Trimmed reads were mapped to a concatenated MRL-CAST genome using STAR v2.5.4b (Dobin et al., 2013), discarding any reads that did not map uniquely to one of the reference genomes (arguments: --outFilterMultimapNmax 1 -- outFilterMultimapScoreRange 1). Requiring that reads map uniquely to one genome ensures we only consider reads overlapping a heterozygous site in F1 individuals. This resulted in an average of 41,215,441, 60,938,290, 76,012,313 uniquely mapped reads for each immune, endothelial, and fibroblast library, respectively. Reads overlapping exonic regions were summed to generate a total count for each gene based on the Ensembl GRCm38 annotation. Cases where reads only mapped to one allele were discarded as they likely reflect SNP calling errors or genomic imprinting. Approximately 50% of reads mapped uniquely to both CAST and MRL, indicating little evidence of mapping bias (see Fig. S8). DESeq2 (Love et al., 2014)(v1.34.0; R computing environment, v4.1.2) was used to perform a variance stabilizing transformation for principal component analysis and perform regularized log2 transformation of the count data (which minimizes differences between samples for rows with small counts and normalizes with respect to library size) for visual comparison of read count data in Fig. 2.

#### Identifying allele-specific expression

DESeq2 was used to identify allele-specific expression and condition-specific ASE (i.e., diffASE) (Love, 2017; Love et al., 2014). Allele-specific expression analyses were restricted to genes with at least 30 reads in each condition (wound type, allele) and non-zero values for >4 alleles across individuals. We analyzed allele-specific reads from each cell population separately with DESeq2 with the model “∼tissue + tissue:sample + tissue:allele” (where “:” denotes an interaction term between two variables in DESeq2). Here, the term “tissue” is the wound site, for differences between ear and dorsal wounds. “Sample” refers to the sample pool the allele-specific sample pertains to, and the term accounts for variation among the different sample pools within wound site groups. “Allele” refers whether allele-specific reads are mapped preferentially to MRL or CAST, and the interaction between allele and tissue is used to estimate the MRL vs CAST allele ratios separately between wound sites. “DiffASE” genes are identified via a contrast between CAST/MRL ratios in ear and dorsal (DESeq2, Wald test). Consequently, significant cases represent scenarios in which the log2 fold change of CAST/MRL differ between wound types. As read counts come from MRL and CAST come from the same sequencing library, library size factor normalization was disabled by setting SizeFactors = 1 (Love, 2017). A false discovery rate correction was applied using the Benjamini-Hochberg method for each comparison. R Code is available on Figshare (https://doi.org/10.6084/m9.figshare.c.6025157).

#### Overlap with previous QTL mapping for ear hole closure

Marker locations and LOD scores for a model with additive and dominance values for ear wound closure are as described by Cheverud et al. 2014 (LOD and marker scores for analysis provided by J. Cheverud). Marker locations were converted from mm9 to mm10 using LiftOver. Autosomal genes were annotated to their closest genetic marker using BEDTools (Quinlan and Hall, 2010) (tool: “closest”) based on Ensembl mm10 gene start and end coordinates. Liftover coordinates of QTL support intervals defined by Cheverud et al. 2014 were used to identify overlap with diffASE genes within QTL using the BEDTools (tool: “intersect”). QTL mapping was performed using LG, the progenitor of MRL. MRL and LG mice share ∼75% of their genome, and both shared and unique QTL from these lines contribute to advanced wound healing (Cheverud et al., 2012, 2014). Consequently, comparisons with this study will be restricted to identifying QTL shared between the lines. However, this should only make our enrichment tests more conservative and for shared causal regions.

#### Enrichment analyses

GO enrichment analyses were performed with PANTHER (Mi et al., 2021), using a foreground list of genes of interest vs. a background list of all genes with sufficient expression to be tested in a cell population (GO Ontology database released 2019-12-09). Mutant phenotype enrichment tests were performed with modPhEA (Weng and Liao, 2017), also using a foreground list of genes and a background list of all genes with sufficient expression to be tested in a cell population.

Enrichment of GO terms and mutant phenotypes for diffASE genes are available in File S1. Enrichment for wound healing terms for fibroblasts was found for diffASE genes at both the FDR<0.05 and FDR<0.1 cut-offs.

#### Fibroblast cell culture

Fibroblast cells were isolated from MRL and CAST dorsal and ear skin for *in vitro* analysis. Following dissection of the dermis from the dorsum and ear, tissue was washed in PBS and finely minced using sterile scissors. Tissue was then digested in collagenase type IV (1500 U/mL in DMEM) at 37 °C, with agitation at 150 rpm, for 1 hour. Enzyme activity was quenched by addition of FBS-enriched media, and digested tissue was successively strained through 300µm followed by 100µm cell strainers. Filtered samples were then centrifuged at 1500 rpm for 5 minutes at 4 °C to obtain a cell pellet. Pelleted cells were resuspended and plated in fibroblast culture media (DMEM + Glutamax media [ThermoFisher, Cat: 10569010] enriched with 10% fetal bovine serum [ThermoFisher, Cat: 10082147] and 1% Antibiotic-Antimycotic [ThermoFisher, Cat: 15240062]) then grown until confluency in tissue culture incubators kept at 37 °C and 5% CO2. Cells used for experiments were between passages 2-4. For *in vitro* analysis of CFH expression, fibroblasts were seeded onto coverslips at 15,000 cells/coverslip for immunostaining (see section below).

#### Immunostaining of cells and wounds

For both OCT wound section slides and cell-seeded coverslips, immunofluorescent staining was performed as follows. Samples were washed twice in Tween 20 (Sigma-Aldrich, St. Louis, MO) followed by one wash in PBS. Samples were then blocked for 1 hour with Power Block (Biogenex, Fremont, CA) prior to addition of anti-CFH primary antibody (LS bio, LS-C819285, 1:200).

Samples were washed then incubated for 1 hour with Alexa Fluor 488 anti-rabbit secondary antibody (Invitrogen, Waltham, MA). Finally, samples were mounted in Fluoromount-G mounting solution with or without DAPI (ThermoFisher Scientific, Waltham, MA). Fluorescent images were acquired with a LSM880 inverted confocal, Airyscan, AiryscanFAST, GaAsP detector upright confocal microscope.

