## Supplemental Figures and Tables for "Allele-specific expression reveals genetic drivers of tissue regeneration in mice"

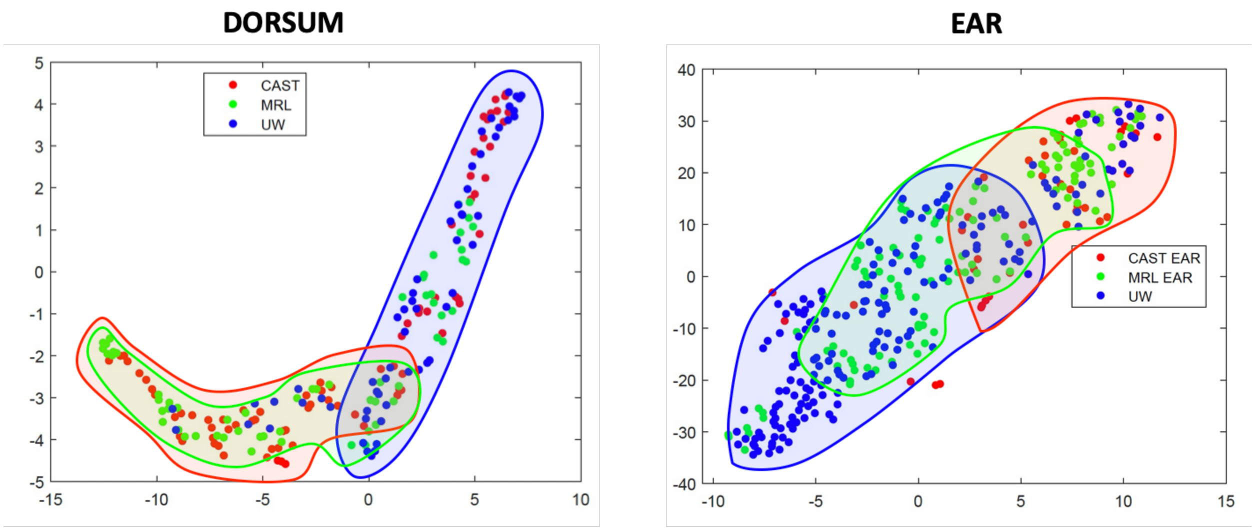


**Figure S1.** T-distributed stochastic neighbor embedding (t-SNE) plots of quantified extracellular matrix (ECM) ultrastructural parameters, based on picrosirius red histology of dorsal (left) and ear (right) wounds, showing overall similarities/differences in ECM ultrastructure between wound and skin conditions. Each dot represents quantified parameters from one histologic image. Overlaid shaded regions highlight clustering of ECM properties by biological condition.

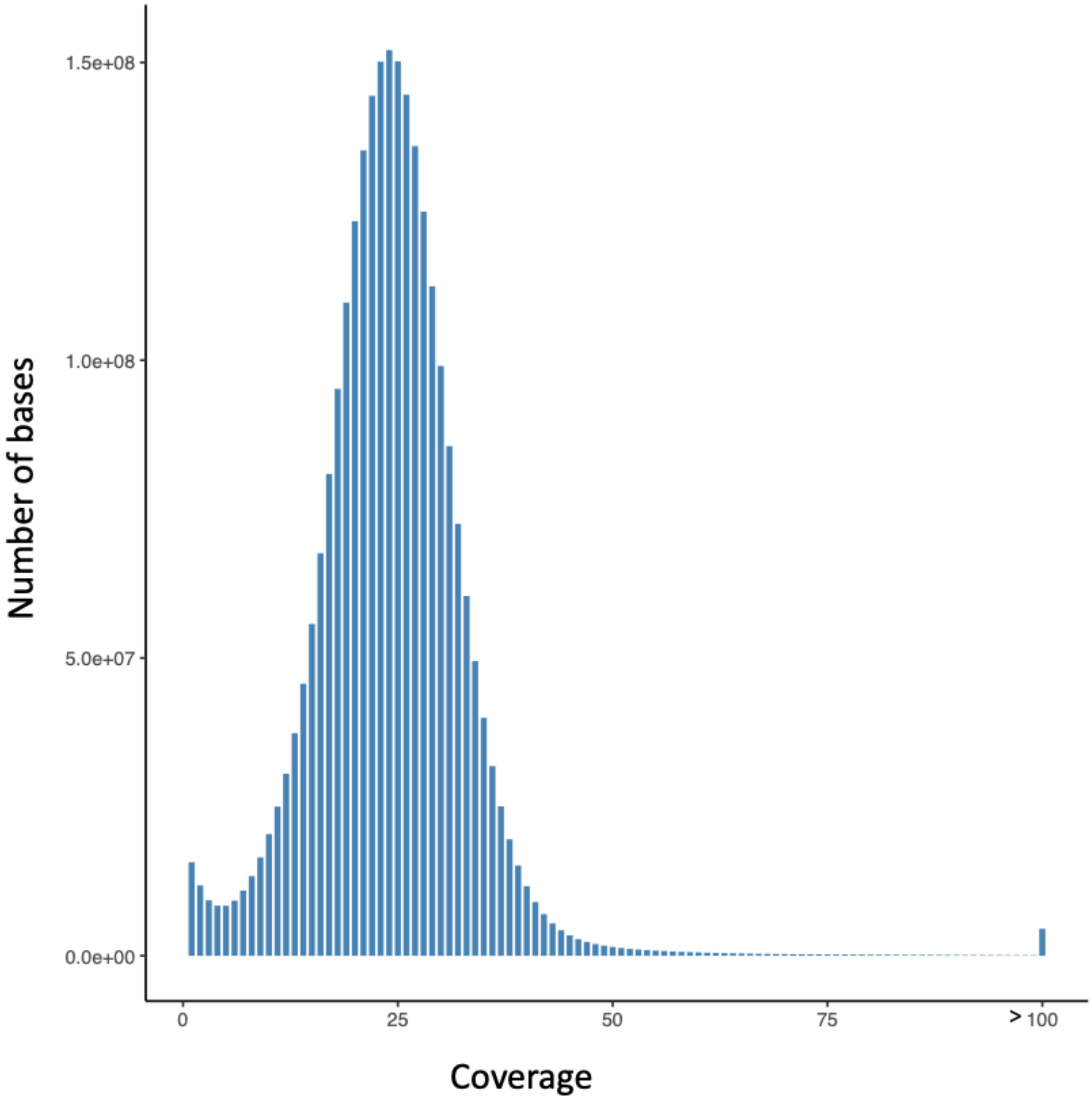


**Figure S2.** Sequencing coverage obtained from MRL genome sequencing. Histogram shows number of bases at each level of coverage.

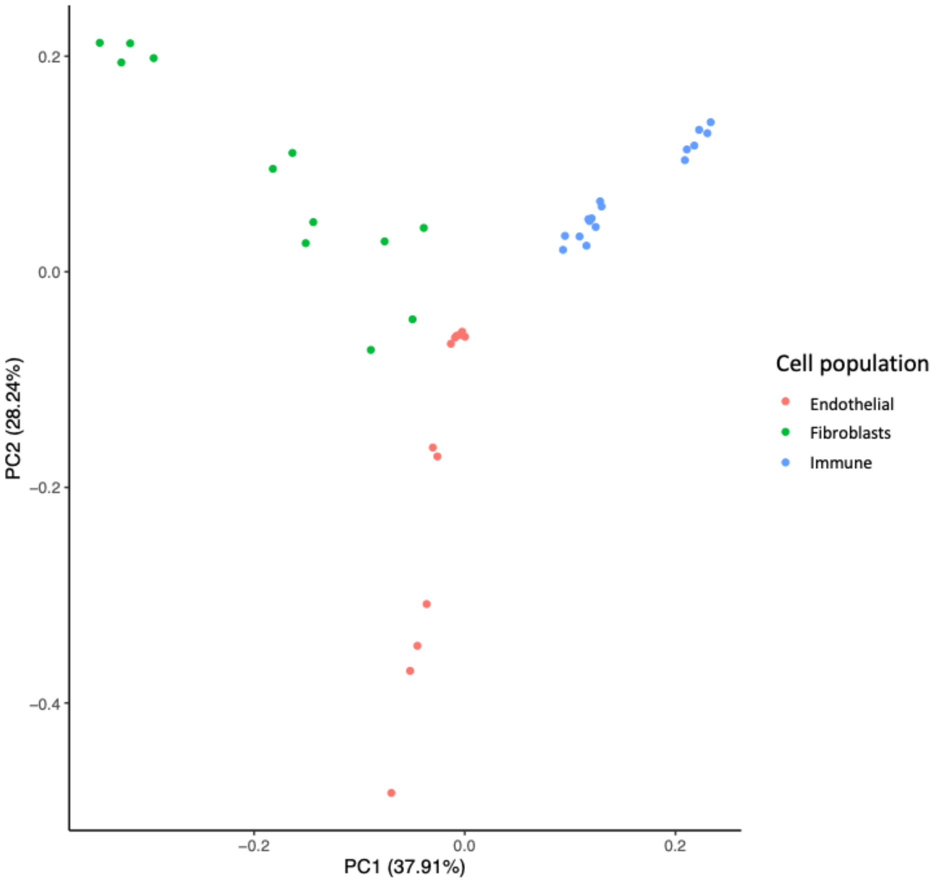


**Figure S3.** Principal component analysis of RNA-seq data clearly separates samples by cell population/type (endothelial cells, fibroblasts, or immune cells).

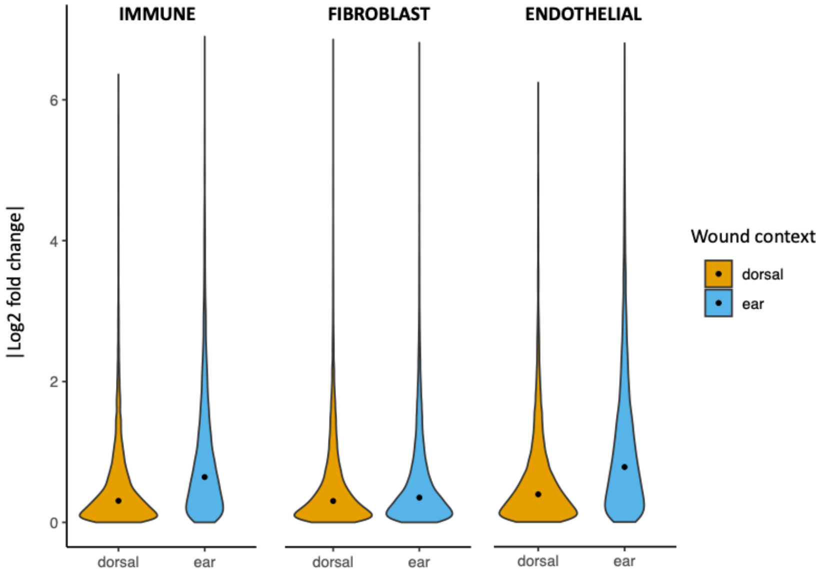


**Figure S4.** Differences in expression between CAST and MRL alleles (i.e., $|\log_2$ fold change$|$) in ear and dorsal wounds in each cell population. Black circles represent medians.

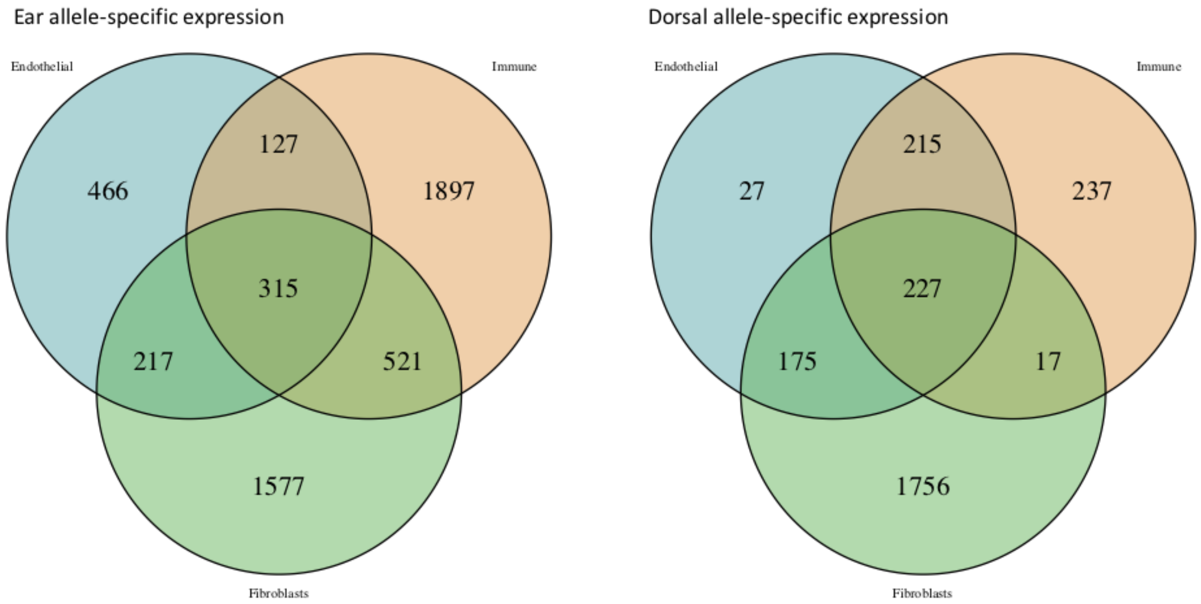


**Figure S5.** Overlap of allele-specific expression (ASE) detected in different cell populations in ear (left) and dorsal (right) wounds, showing number of genes with significant ASE (FDR < 0.05 for MRL vs. CAST allelic expression) in each individual cell type or in multiple cell types (overlapping regions).

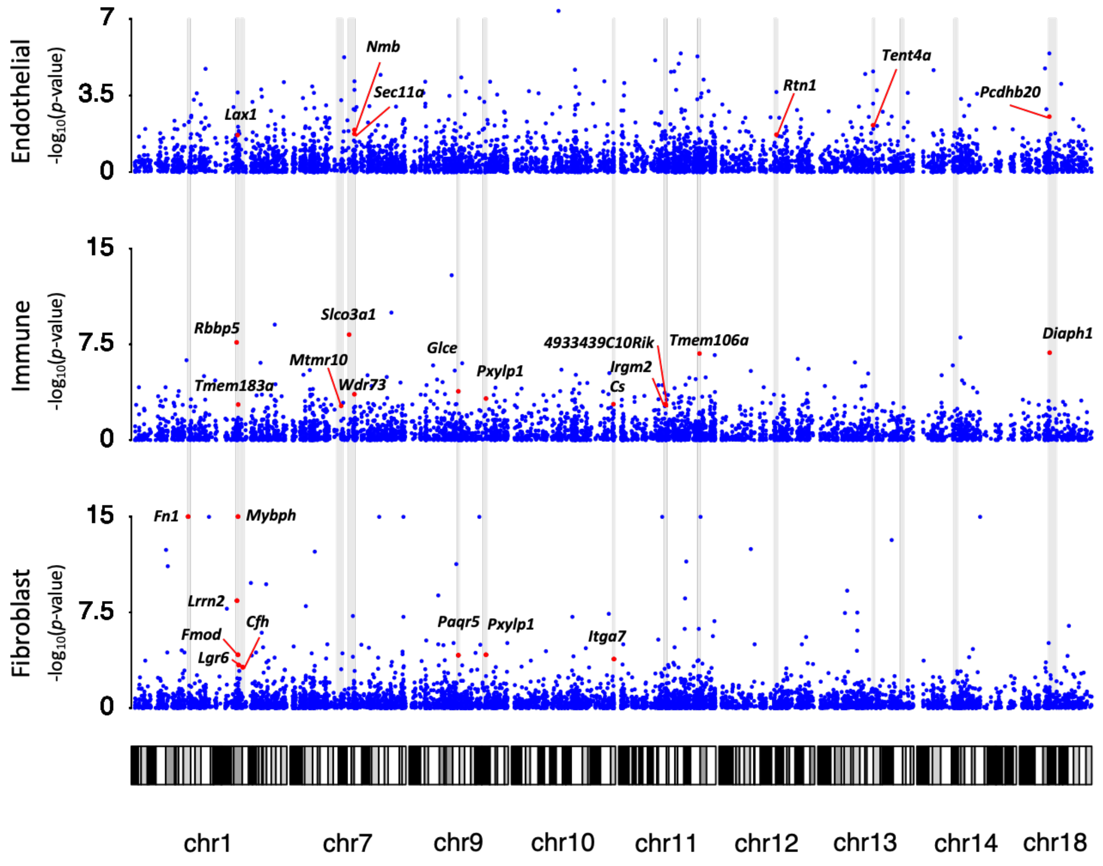


**Figure S6.** Genes with diffASE overlapping 19 QTL support intervals The -log($p$-value) for diffASE for each gene (red and blue dots) are plotted versus genomic position for each cell type. Gray regions highlight QTL support intervals for which we identified a gene with significant diffASE. Red dots highlight genes with significant diffASE within a QTL interval.

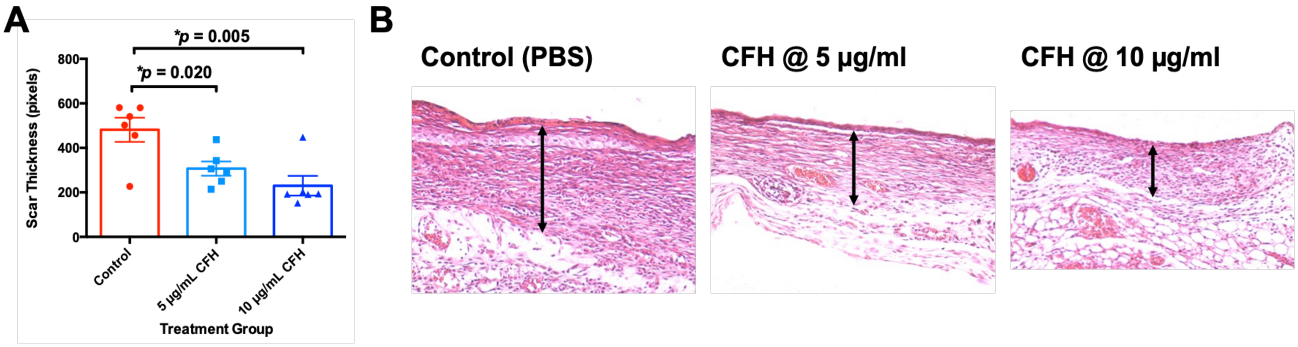


**Figure S7:** Quantified scar thickness (**A**) and representative H&E histology (**B**) of wildtype mouse wounds treated with PBS (control) or CFH at varying doses (5 or 10 µg/mL; see Methods for full details/dosing).

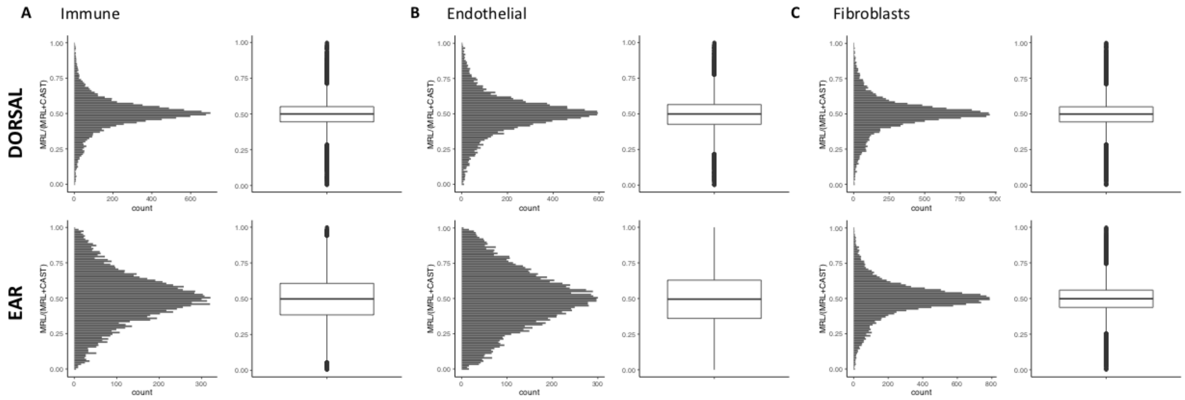


**Figure S8.** Allelic proportions (MRL reads/CAST reads per gene, summed across replicates) in (**A**) immune cells, (**B**) endothelial cells, and (**C**) fibroblasts. Allelic proportions center on 0.5 in each condition.

**Supplemental Tables:**

**Table S1:** Reads mapped per library

| Cell type | Wound Type | Mouse/Pool | ID | Total reads | Uniquely mapped[1] |
|---|---|---|---|---|---|
| Immune | Ear | 1 | immune1 | 107291479 | 38083153 |
| Immune | Dorsal | 1 | immune2 | 113414580 | 41345663 |
| Immune | Ear | 2 | immune3 | 126081376 | 44199848 |
| Immune | Dorsal | 2 | immune4 | 121655577 | 43424110 |
| Immune | Ear | 3 | immune5 | 103893903 | 34075181 |
| Immune | Dorsal | 3 | immune6 | 108072579 | 36835581 |
| Immune | Ear | 4 | immune7 | 137991403 | 44094860 |
| Immune | Dorsal | 4 | immune8 | 131912247 | 47665128 |
| Endothelial | Dorsal | 5 | endo1 | 236886813 | 71816524 |
| Endothelial | Ear | 5 | endo2 | 253802304 | 72722333 |
| Endothelial | Dorsal | 6 | endo3 | 193057852 | 50413225 |
| Endothelial | Ear | 6 | endo4 | 199289775 | 53015720 |
| Endothelial | Dorsal | 7 | endo5 | 216283212 | 59504234 |
| Endothelial | Ear | 7 | endo6 | 237009122 | 58157705 |
| Fibroblasts | Dorsal | 8 | fibro1 | 235618531 | 80417330 |
| Fibroblasts | Ear | 8 | fibro2 | 229547131 | 86880172 |
| Fibroblasts | Dorsal | 9 | fibro3 | 230492907 | 64244207 |
| Fibroblasts | Ear | 9 | fibro4 | 219631076 | 69345695 |
| Fibroblasts | Dorsal | 10 | fibro5 | 244019067 | 72484120 |
| Fibroblasts | Ear | 10 | fibro6 | 240201101 | 82702356 |

[1]Mapped uniquely to either CAST or MRL

**Table S2:** The number of genes with sufficient read counts (30 reads per wound type, allele) to be analyzed for allele-specific expression in each cell type

| Cell types | Number of genes with sufficient coverage |
|---:|---:|
| Immune | 10,555 |
| Endothelial | 11,831 |
| Fibroblast | 14,674 |

**Table S3:** ASE results from DESeq2 at FDR<0.1 and FDR<0.05.

| Cell types, condition | FDR<0.1 | FDR<0.05 |
|---|---|---|
| Ear, immune | 3493 | 2860 |
| Dorsal, immune | 813 | 696 |
| Ear vs. Dorsal, immune | 705 | 432 |
| Ear, endothelial | 1496 | 1125 |
| Dorsal, endothelial | 754 | 644 |
| Ear vs. Dorsal, endothelial | 141 | 91 |
| Ear, fibroblasts | 3270 | 2630 |
| Dorsal, fibroblasts | 2668 | 2175 |
| Ear vs. Dorsal, fibroblasts | 319 | 235 |

**Table S4:** Directionality and magnitude for genes with allele-specific expression in both ear and dorsal wounds.

| Cell population | Fold change in same direction in ear and dorsal wounds | Significantly different magnitude (FDR<0.05) | Significantly different magnitude (FDR<0.1) |
|---|---|---|---|
| Endothelial | 371/371 | 1 | 2 |
| Fibroblast | 1360/1377 | 51 | 59 |
| Immune | 568/568 | 25 | 36 |

**Table S5:** Genes with differential allele-specific expression in fibroblasts associated with mutant phenotypes related to abnormal response to injury (annotated to mutant phenotypes based on ModPhEA).

| Gene name | Mutant phenotypes associated with abnormal response to injury | Greater | Up-regulation in Ear | Up-regulation in Dorsal |
|---|---|---|---|---|
| *C3* | decreased susceptibility to injury, accelerated wound healing | Ear | CAST | MRL |
| *Hspb1* | abnormal wound healing, delayed wound healing, impaired wound healing | Ear | MRL | MRL |
| *Cav1* | abnormal vascular wound healing, impaired wound healing, cardiac fibrosis | Ear | CAST | CAST |
| *Fn1* | impaired wound healing | Ear | CAST | MRL |
| *Aebp1* | impaired wound healing | Ear | CAST | CAST |
| *Penk* | delayed wound healing | Dorsal | MRL | CAST |
| *Slpi* | abnormal wound healing, delayed wound healing, impaired wound healing | Ear | MRL | No ASE |
| *Spp1* | abnormal vascular wound healing, abnormal wound healing, altered response to myocardial infarction, delayed wound healing, increased susceptibility to injury | Ear | MRL | No ASE |
| *Cp* | abnormal response to injury | Dorsal | No ASE | MRL |
| *Gpx3* | decreased bleeding time, abnormal thrombosis, increased platelet aggregation, decreased susceptibility to ischemic brain injury | Ear | CAST | No ASE |
| *Thbs4* | delayed wound healing; cardiac fibrosis | Ear | MRL | MRL |
| *Anxa1* | delayed wound healing | Ear | CAST | No ASE |
| *F3* | abnormal vascular wound healing, abnormal blood coagulation, abnormal thrombosis | Ear | CAST | No ASE |
| *Comp* | abnormal vascular wound healing | Dorsal | No ASE | CAST |
| *Pla2g5* | altered response to myocardial infarction | Dorsal | No ASE | CAST |
| *Adora2b* | increased vascular permeability, increased susceptibility to injury, decreased susceptibility to injury | Ear | CAST | No ASE |

| | | | | |
|---|---|---|---|---|
| *Plec* | abnormal wound healing, impaired wound healing, skeletal muscle fiber necrosis | Dorsal | No ASE | MRL |
| *Ccn1* | increased susceptibility to injury | Ear | CAST | No ASE |
| *Nt5e* | altered response to myocardial infarction, decreased bleeding time, abnormal thrombosis | Ear | MRL | CAST |
| *Rcan1* | altered response to myocardial infarction | Dorsal | No ASE | CAST |
| *Il1rn* | abnormal vascular wound healing | Ear | MRL | No ASE |
| *Clic4* | delayed wound healing, abnormal corneal wound healing | Ear | MRL | No ASE |
| *Gfap* | increased susceptibility to injury | Ear | CAST | CAST |
| *Cst3* | decreased susceptibility to ischemic brain injury | Ear | CAST | MRL |
| *Mmp2* | altered response to myocardial infarction | Ear | CAST | CAST |
| *F13a1* | abnormal blood coagulation, hemorrhage, increased bleeding time | Dorsal | No ASE | MRL |

**Table S6:** Genes associated with differential allele-specific expression in fibroblasts associated with the GO terms "response to wounding" and "wound healing" (annotated with PANTHER, GO Ontology database released 2019-12-09)

| Gene name | Function | Greater | Up-regulation in Ear | Up-regulation in Dorsal |
|---|---|---|---|---|
| *Cyr61/Ccn1* | growth factor | Ear | CAST | No ASE |
| *C3* | complement component; cytokine; serine protease inhibitor | Ear | CAST | MRL |
| *Tspan32* | | Dorsal | No ASE | CAST |
| *Lgr6* | extracellular matrix protein; receptor | Ear | CAST | No ASE |
| *Slc7a11* | | Ear | CAST | No ASE |
| *Plec* | intermediate filament binding protein | Dorsal | No ASE | MRL |
| *Ninj1* | cell adhesion molecule | Dorsal | MRL | CAST |
| *Cfh* | | Ear | MRL | No ASE |
| *Anxa1* | | Ear | CAST | No ASE |
| *Cav1* | G-protein modulator; membrane traffic protein; structural protein; transmembrane receptor regulatory/adaptor protein | Ear | CAST | CAST |
| *Jaml* | cell adhesion molecule; voltage-gated sodium channel | Ear | CAST | No ASE |
| *Adrb2* | G-protein coupled receptor | Ear | CAST | No ASE |
| *Mymk* | cell adhesion molecule | Ear | CAST | No ASE |
| *Fn1* | signaling molecule | Ear | CAST | MRL |
| *F13a1* | acyltransferase | Dorsal | No ASE | MRL |
| *Comp* | | Dorsal | No ASE | CAST |
| *Fgf7* | growth factor | Ear | CAST | No ASE |

| | | | | |
|---|---|---|---|---|
| *Ecrg4* | | Dorsal | CAST | No ASE |
| *Mmp2* | metalloprotease | Ear | CAST | CAST |
| *Sulf2* | hydrolase | Dorsal | No ASE | CAST |
| *Gfap* | | Ear | CAST | CAST |
| *Nrep* | | Ear | CAST | No ASE |
